# WikiGOA: Gene set enrichment analysis based on Wikipedia and the Gene Ontology

**DOI:** 10.1101/2022.09.15.508149

**Authors:** Tiago Lubiana, Thomaz Lüscher Dias, Débora Guerra Peixe, Helder Takashi Imoto Nakaya

## Abstract

Gene sets curated to Gene Ontology terms are widely used by the transcriptomics community
Presence in Wikipedia is a common proxy for the relevance of a concept.
In this work, we describe the use of Wikidata to generate a dataset comprising only gene sets with a corresponding Wikipedia page.
We refer to the dataset as “WikiGOA”, standing for “Wikipedia Gene Ontology Annotations”
We use the dataset to analyze gene expression data and show that it provides readily understandable results.
We envision WikiGOA to be useful for exploring complex biological datasets both in academic research and educational contexts.

**Note:** This report was written in a non-standard, experimental format, where assertions are expressed in bullet points. This was done to clarify statements and assumptions, simplify reading and pave the way for conversion to structured formats (e.g., nanopublications). [1]

## Introduction

- The analysis of omics data relies on integrating datasets with curated databases of gene function annotation.
- Gene sets curated to Gene Ontology (GO) terms are widely used by the transcriptomics community for Over Representation Analysis and Gene Set Enrichment Analysis.[2–4]
- A large number of gene sets is a double-edged sword: there is a tradeoff between coverage of niche fields and the quick interpretability of results.
- Furthermore, the statistical power of the analysis is reduced when too many comparisons are present.
- One challenge is identifying which GO terms are the most relevant for the task.
- Wikipedia is a common lay metric of relevance, as the creation of pages is bound by notability criteria (https://en.wikipedia.org/wiki/Wikipedia:Notability).
- Wikidata is a Linked Open Data hub providing links to GO identifiers and Wikipedia.[5,6]
- In this work, we describe the use of Wikidata to generate a dataset comprising only gene sets for Gene Ontology Biological Processes with a corresponding Wikipedia page.
- We refer to the datasets as “WikiGOA”, standing for “Wikipedia Gene Ontology Annotations”
- We use the dataset to analyze gene expression data and show that it provides readily understandable results, useful for exploring complex biological datasets.
- A visual representation of our workflow is shown in **Figure** 1.

**Figure 1:**
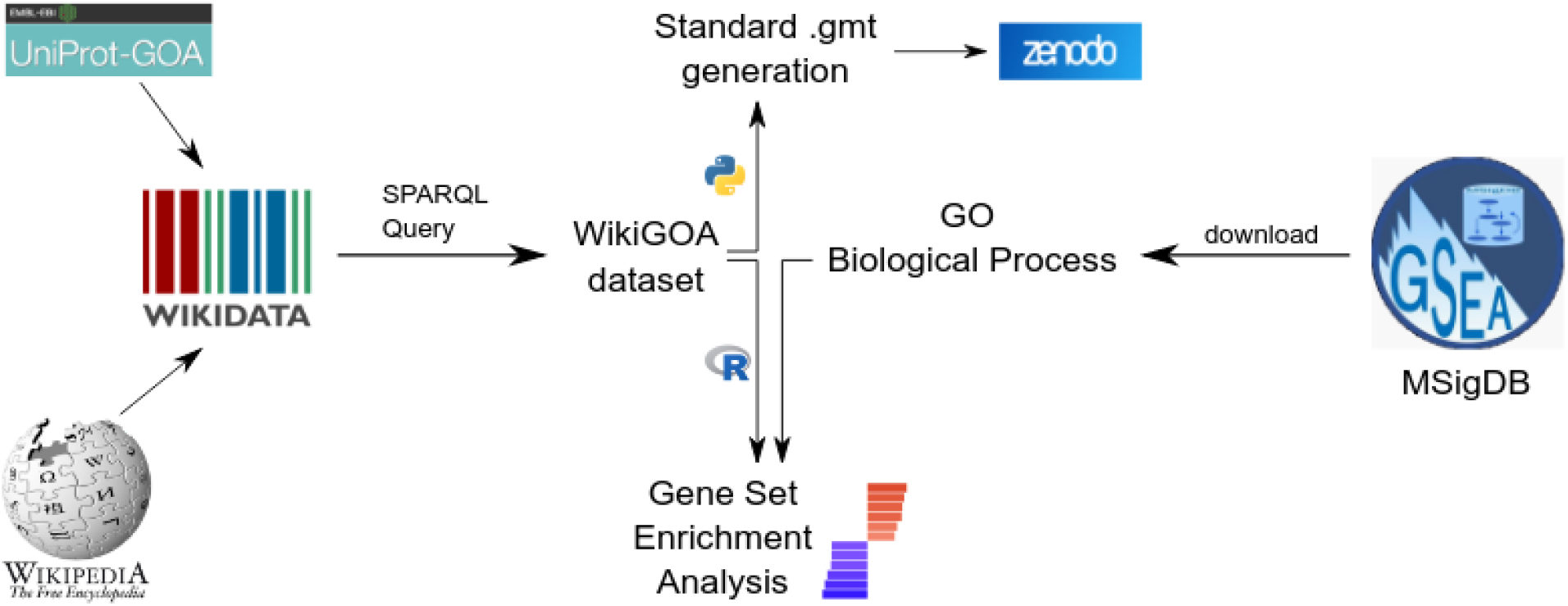
The WikiGOA dataset. Using SPARQL, we extracted gene sets from Wikidata (reconciled mostly from UniProt-GOA). Only GO terms with associated English Wikipedia pages were selected. We rendered datasets in standard formats and compared the performance with default Gene Ontology annotations from MSigDB.

## Methods

### Dataset preparation

- The dataset for Biological Process Gene Sets, released under the CC-BY-4.0 License, was downloaded from MSigDB (http://www.gsea-msigdb.org/gsea/downloads.jsp) .[7][8]
- Gene Ontology terms with English Wikipedia pages were obtained from Wikidata using the SPARQL query language via the Wikidata Query Service:

~~~
SELECT DISTINCT ?site ?item ?itemLabel ?go
 ?gene_symbol ?entrez ?ensembl_gene_id .
WHERE
{
 ?item wdt:P686 ?go .
 ?protein wdt:P682 ?item ;
       wdt:P703 wd:Q15978631 ;
       wdt:P702 ?gene .
 ?gene wdt:P353 ?gene_symbol ;
       wdt:P351 ?entrez ;
       wdt:P594 ?ensembl_gene_id .
 ?sitelink schema:about ?item ;
       schema:isPartOf <https://en.wikipedia.org/> .
 ?item rdfs:label ?itemLabel. FILTER (LANG (?itemLabel) = “en”)
}
~~~

### Code Snippet 1: SPARQL query to generate the Wikipedia-based gene sets

https://w.wiki/5DsS

- The dataset was prepared via Wikidata and includes HGNC gene symbols, Ensembl IDs and Entrez IDs
- The final dataset had 269 gene sets, comprising all Gene Ontology Biological Process terms with Wikidata cross references to Wikidata as of June 2022.
- The datasets for enrichment analysis are available in CC0 both as a .tsv table and in the standard .gmt format at https://doi.org/10.5281/zenodo.6624354.
- Gene Ontology Annotations were obtained from Wikidata, where most of the protein annotations come from the UniProt-GoA database [9] via mappings done by the ProteinBoxBot [6].
- The list of all Gene Ontology terms with Wikipedia pages in all categories (Biological Process, Cell Component, and Molecular Function) is also available via the Wikidata Query Service: https://w.wiki/5Dsh

## Results

- We qualitatively compare the performance of WikiGOA and the MSigDB Gene Ontology Biological Processes dataset (MSigDB GOBP) for two publicly available gene expression datasets; one about SARS-CoV-2 infected cells [10] and the other about post-mortem samples of schizophrenia patients. [11] (**Figure 2**)

**Figure 2:**
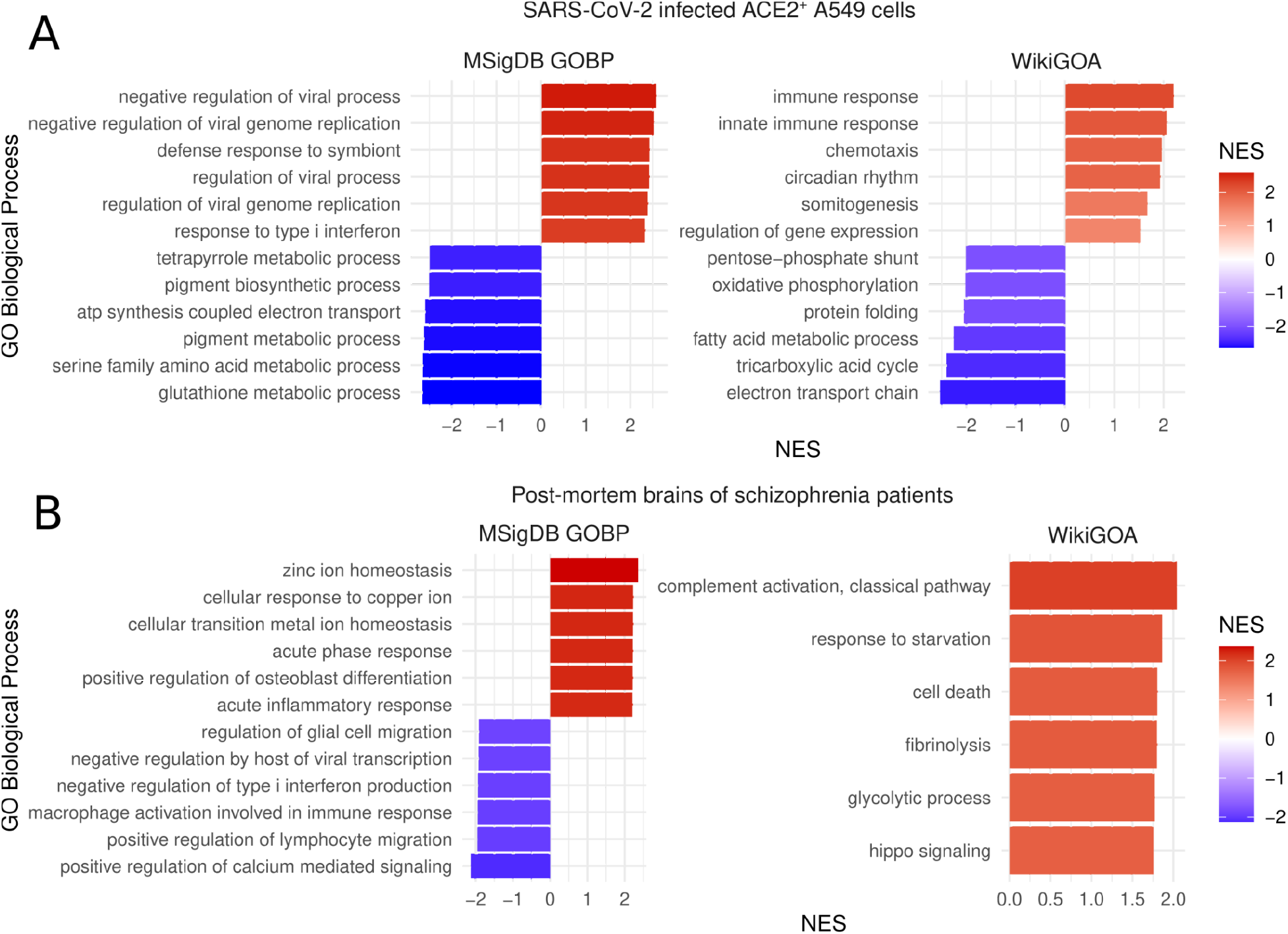
Comparison of the performance of WikiGOA and MSigDB datasets. Top 6 up positively and negatively enriched biological process for both WikiGOA and MSigDB GOBP for A) ACE+ A549 cells infected with SARS-CoV-2 in comparison to non-infected cells and B) brains from schizophrenia patients in comparison to non-schizophrenia brains.
- The gene set names of WikiGOA are different from MSigDB, as WikiGOA uses the Wikidata labels and MSigDB uses the original Gene Ontology labels.
- Gene sets are different in composition, as WikiGOA uses the gene annotations on Wikidata. It is unclear which version of GO annotations are used in MSigDB (more information in gsea-msigdb.org/gsea/msigdb/human/collection_details.jsp#GO)
- The WikiGOA dataset contains gene sets corresponding to 269 different Gene Ontology terms, while the MSigDB dataset contains 7658.

### SARS-CoV-2 dataset

- The SARS-CoV-2 dataset compared the transcriptomes of ACE+ A549 cells (derived from a lung carcinoma) that were infected with SARS-CoV-2 to identical cells that were not infected.
- For that dataset, WikiGOA highlights an upregulation of immune response genes and a general downregulation of the basal energy metabolism of the cells, consistent with what is known in the literature about the biology of SARS-CoV-2. [12,13]
- The gene sets selected by using WikiGoA that could be matched to gene sets in MSigDB displayed lower adjusted p-values for WikiGOA (**Table 1**).

**Table 1.** Adjusted p-values for FGSEA sets for the COVID-19 dataset
- This is expected, given that the multiple tests correction was applied to a lower amount of sets. Note that this may not always be the case, as the gene composition of the sets is different between the datasets.

| Pathway name | PAdj - WikiGOA | PAdj - MSigDB | Size - WikiGOA | Size - MSigDB |
| --- | --- | --- | --- | --- |
| circadian rhythm | 0.081 | 0.082 | 68 | 147 |
| immune response | 0.081 | 0.082 | 168 | 908 |
| innate immune response | 0.081 | 0.082 | 275 | 456 |
| electron transport chain | 0.092 | 0.117 | 64 | 138 |
| muscle contraction | 0.092 | 0.775 | 55 | 178 |
| oxidative phosphorylation | 0.092 | 0.113 | 12 | 125 |
| tricarboxylic acid cycle | 0.092 | 0.095 | 30 | 28 |
| fatty acid metabolic process | 0.094 | 0.153 | 130 | 261 |
| somitogenesis | 0.094 | 0.632 | 38 | 39 |
| protein folding | 0.097 | 0.126 | 146 | 171 |

### Schizophrenia dataset

- The schizophrenia dataset compared the post-mortem brains of people diagnosed with schizophrenia with the post-mortem brains of non-schizophrenia matched controls.
- For that dataset, WikiGOA highlights a series of different upregulated processes, including the complement pathway and hippo signaling pathway, both of which have been associated with schizophrenia before. [14] [15]
- The upregulation of fibrinolysis is opposite to previous studies that found a global hipofibrinolysis and hypercoagulability in schizophrenia patients. [16] We reason that the difference is welcome, as it shows WikiGOA as capable of raising unexpected situations that may lead, in practice, to novel interpretations and explanations of biological phenomena.
- The same trend of lower adjusted P-values seen for the COVID-19 dataset was also seen for some pathways in the Schizophrenia dataset, although not all. (**Table 2**) This is due to the difference in the composition of the gene sets, noticeable in their size.

**Table 2.** Adjusted p-values for FGSEA sets for the Schizophrenia dataset

| Pathway name | PAdj - WikiGOA | PAdj - MSigDB | Size - WikiGOA | Size - MSigDB |
| --- | --- | --- | --- | --- |
| immune response | 0.08 | 0.052 | 270 | 1187 |
| innate immune response | 0.08 | 0.052 | 367 | 578 |
| epithelial to mesenchymal transition | 0.092 | 0.052 | 36 | 139 |
| fibrinolysis | 0.092 | 0.107 | 13 | 16 |
| response to starvation | 0.092 | 0.052 | 31 | 168 |
| carbohydrate metabolic process | 0.099 | 0.19 | 204 | 509 |
| hippo signaling | 0.099 | 0.107 | 26 | 39 |
| wound healing | 0.099 | 0.052 | 81 | 359 |

## Discussion

- In summary, for both datasets, the WikiGOA enrichment highlighted simple terms, that are easier to understand.
- It also seems that the terms picked by WikiGOA are less redundant than the ones retrieved in the GOBP analysis.
- These 2 characteristics (readability and independence of the terms retrieved) are arguably more useful for a first-pass overview of the data than the fine-grained results brought when using MSigDB.
- The lower complexity of the retrieved terms and the guaranteed presence of the concepts on Wikipedia also arguably present a more friendly environment for bioinformatics students learning to handle gene sets.
- One limitation is that, by design, WikiGOA will have a gross granularity, as only major biological processes have Wikipedia pages. Thus, for bioinformatics research, we recommend using WikiGOA alongside fine-grained datasets, such as MSigDB, for a more complete overview of the data at hand.

## Conclusion

- The Wikipedia Gene Ontology Annotations dataset is a simple but powerful addition to the arsenal of the transcriptomics community.
- We envision its usage for quick exploration of gene expression data both in the context of bioinformatics research and higher education.

## Data analysis

- The gene sets were analyzed in R using the fGSEA package (https://www.biorxiv.org/content/10.1101/060012v3)
- Code is available at https://github.com/luxeredias/WikiGOA.

## Funding

- TL is supported by a grant from the São Paulo Research Foundation (#19/26284-1).
- TLD is supported by a grant from the Eboplus consortium.

## Notes

### Competing Interest Statement

The authors have declared no competing interest.

